## Supplemental information for "SDE2 Integrates into the TIMELESS-TIPIN Complex to Protect Stalled Replication Forks"

**Table S1. Oligonucleotides**

| **siRNA target sequences** | | |
|---|---|---|
| SDE2-1 aaACGGCAATGGCCTACTAAA | Qiagen | custom |
| SDE2-5 acGCAGTTATTGATAAGGAAA | Qiagen | custom |
| SDE2-7 ctGAATAAGGATAAAGAGACA | Qiagen | custom |
| TIM-1 ggGTAGCTTAGTCCTTTCAAA | Qiagen | custom |
| TIM-2 aaGAGCTAAGAAGCCTAGGGG | Qiagen | custom |
| TIPIN cgTGATTGACCTACCAGATTA | Qiagen | custom |
| FANCD2 ttGGAGGAGATTGATGGTCTA | Qiagen | custom |
| ETAA1 atTGACAAAGCAGTTAGGTAA | Ambion | Cat# s29018 |
| TOPBP1 aaCTCACCTTATTGCAGGAGA | Ambion | Cat# s21823 |
| SMARCAL1 caGCTTTGACCTTCTTAGCAA | Qiagen | custom |
| MRE11 atGAAAGGCTCTATCGAATGT | Qiagen | custom |
| CtIP aaCGAATCTTAGATGCACAAA | Qiagen | custom |
| DNA2 gcATAGCCAGTAGTATTCGAT | Qiagen | custom |
| EXO1 ttGCCTGAGAATAATATGTCT | Qiagen | custom |
| BRCA2 ttGAAGAATGCAGGTTTAATA | Qiagen | custom |

| **SDM Primers** | |
|---|---|
| SDE2 Δ108-150 | CGGGATCTCAGTGGAAGGTTCACCAGCCCC GACTAC |
| SDE2 K132A/K135A | GAG CGA GAG GCT GAA gcG GAG CAG gcG CGG CTG GAG CGA CT |
| TIM Δ1132-1208 | AAA GAG CAC CGA GCA tAA GCC CTG AGG GCC |
| TIM Δ882-1208 | C AAG GAC TTC CAA AGG tAA GGA ACC CAT ATT G |
| TIM T1078D | GCC TCT GGG CAG GAA gaC TTC TGG CGA ATT CC |
| TIM E1049Q | TTG GTG CCA CTC ACA cAG GAA AAT GAG GAA G |
| TIM E1056Q | AAT GAG GAA GCC ATG cAA AAC GAA CAG TTT C |

**Table S2. Chemicals and Reagents**

| | | |
|---|---|---|
| Hydroxyurea (HU) | Sigma-Aldrich | Cat# H8627 |
| 5-iodo-2´-deoxyuridine (IdU) | Sigma-Aldrich | Cat# I7125 |
| 5-chloro-2´-deoxyuridine (CldU) | Sigma-Aldrich | Cat# C6891 |
| 5-ethynyl-2´-deoxyuridine (EdU) | Thermo Fisher Scientific | Cat# A10044 |
| 5-bromo-2´-deoxyuridine (BrdU) | Sigma-Aldrich | Cat# B5002 |
| $H_2O_2$ | Sigma-Aldrich | Cat# H1009 |
| RNAiMAX transfection reagent | Thermo Fisher Scientific | Cat# 13778150 |
| Genejuice transfection reagent | MilliporeSigma | Cat# 70967 |
| Xfect transfection reagent | Clontech Laboratories | Cat# 631317 |
| cOmplete, EDTA-free protease inhibitor cocktail | MilliporeSigma | Cat# 11873580001 |
| Halt phosphatase inhibitor cocktail | Thermo Fisher Scientific | Cat# 78420 |
| MG132 | Sigma-Aldrich | Cat# C2211 |
| Cycloheximide | Sigma-Aldrich | Cat# C4859 |
| Camptothecin | Sigma-Aldrich | Cat# C9911 |
| Doxycycline hyclate | Sigma-Aldrich | Cat# D9891 |
| Mirin | Sigma-Aldrich | Cat# M9948 |
| Puromycin | Sigma-Aldrich | Cat# P8833 |
| Biotin-phenol (BP) | R&D Systems | Cat# 6241 |
| Thymidine | Sigma-Aldrich | Cat# T1895 |
| Streptavidin, Alexa Fluor 594 | Thermo Fisher Scientific | Cat# S11227 |
| Streptavidin-HRP | Thermo Fisher Scientific | Cat# S911 |
| Streptavidin agarose | Thermo Fisher Scientific | Cat# 20359 |
| Streptavidin agarose | MilliporeSigma | Cat# 69203 |
| Biotin azide | Thermo Fisher Scientific | Cat# B10184 |
| Nocodazole | Sigma-Aldrich | Cat# M1404 |
| Sodium ascorbate | VWR | Cat# 95035-692/S1349 |
| Trolox | Sigma-Aldrich | Cat# 238813 |
| Sodium azide | VWR | Cat# AA14314-22/14314 |
| Aprotinin | Sigma-Aldrich | Cat# A6279 |
| Leupeptin | Sigma-Aldrich | Cat# L2884 |
| Ponceau S | Boston Bioproducts | Cat# ST-180 |
| FLAG M2 affinity gel | Sigma-Aldrich | Cat# A2220 |
| Glutathione agarose | Thermo Fisher Scientific | Cat# 16100 |
| Dynabeads protein G | Thermo Fisher Scientific | Cat# 10003D |
| SYBR Gold nucleic acid gel stain | Thermo Fisher Scientific | Cat# S11494 |

**Table S3. Antibodies**

| | | |
|---|---|---|
| BRCA2 (Ab-1) | MilliporeSigma | Cat# OP-95 |
| CHK1 | Santa Cruz | Cat# sc-8408 |
| pCHK1 S345 | Cell Signaling Technology | Cat# 2341 |
| FANCD2 (FI-17) | Santa Cruz | Cat# sc-20022 |
| FLAG | Sigma-Aldrich | Cat# F1804 |
| GFP (B-2) | Santa Cruz Biotechnology | Cat# sc-9996 |
| GFP (polyclonal) | Abcam | Cat# ab290 |
| γH2AX S139 | Cell Signaling Technology | Cat# 2577 |
| HA (6E2) | Cell Signaling Technology | Cat# 2367 |
| Histone H3 | Abcam | Cat# ab1791 |
| HSC70 (B6) | Santa Cruz | Cat# sc-7298 |
| MCL-1 | Bethyl Laboratories | Cat# A302-715A |
| MCM6 (H-8) | Santa Cruz | Cat# sc-393618 |
| Myc (9E10) | Santa Cruz | Cat# sc-40 |
| ORC-2 | BD Biosciences | Cat# 551178 |
| p97 | Cell Signaling Technology | Cat# 2648 |
| PARP1 | Bethyl Laboratories | Cat# A301-376A-T |
| PARP1 (F-2) | Santa Cruz | Cat# sc-8007 |
| PCNA (PC-10) | Santa Cruz | Cat# sc-56 |
| RPA32 | MilliporeSigma | Cat# MABE285 |
| pRPA32 S4/S8 | Bethyl Laboratories | Cat# A300-245A-M |
| pRPA32 S33 | Bethyl Laboratories | Cat# A300-246A |
| SDE2 | Sigma Atlas | Cat# HPA031255 |
| SMARCAL1 | Santa Cruz | Cat# sc-376377 |
| TIMELESS | Bethyl Laboratories | A300-961A-M |
| TIPIN | Bethyl Laboratories | Cat# A301-474A |
| γ-Tubulin | Bethyl Laboratories | Cat# A302-631A |
| α-Tubulin | Santa Cruz | Cat# sc-32293 |
| Cyclin E (HE12) | Santa Cruz | Cat# sc-247 |
| Cyclin A (B-8) | Santa Cruz | Cat# sc-271682 |
| KU80 | Cell Signaling Technology | Cat# 2753 |
| β-Actin | Thermo Fisher Scientific | Cat# MA5-15739 |
| BrdU (BU-1) | Thermo Fisher Scientific | Cat# MA3-071 |
| BrdU (IdU) (B44) | BD Biosciences | Cat# 347580 |
| BrdU (CldU) (BUI/75 ICR1) | Abcam | Cat# ab6326 |
| Biotin (mouse) | Jackson ImmunoResearch | Cat# 200-002-211 |
| Biotin (rabbit) | Bethyl Laboratories | Cat# A150-109A |
| Trueblot Ultra: anti-mouse IgG HRP | Rockland | Cat# 18-8817-33 |
| Trueblot Ultra: anti-rabbit IgG HRP | Rockland | Cat# 18-8816-33 |
| Normal Rabbit IgG | Millipore-Sigma | Cat# 12-370 |

Figure S1

A

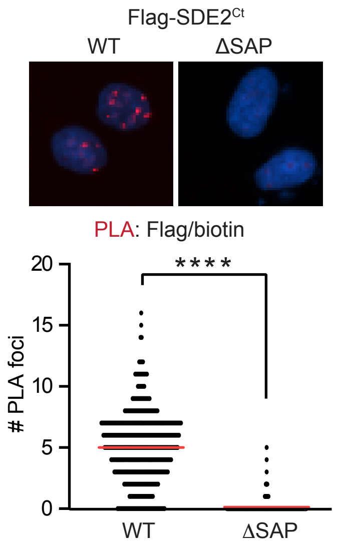


B

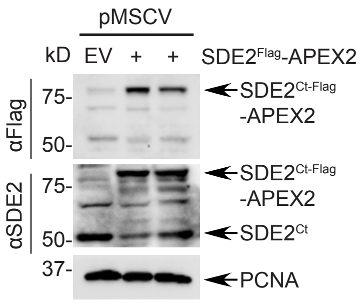


C

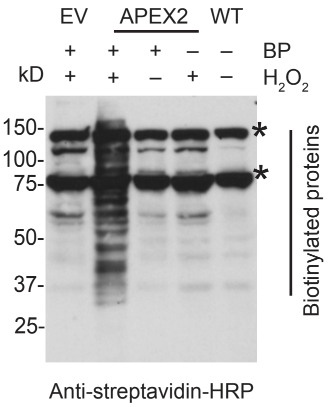

**Supplementary Figure S1 (related to Figure 1).**

**(A)** Top: representative images of PLA foci between Flag-SDE2$^{Ct}$ and EdU. EdU-incubated U2OS cells expressing N-terminal UBL-deleted Flag-SDE2 wild-type (WT) or ΔSAP mutant (aa385-451) were fixed, and PLA foci were visualized by anti-Flag and anti-biotin antibodies. Bottom: dot plot showing PLA foci numbers within Flag:biotin PLA positive cells. Red bars represent the mean (n=2, a representative experiment is shown, ****$P$<0.0001, Mann-Whitney test). **(B)** Western blot (WB) analysis of U2OS cells stably expressing SDE2-APEX2 by retroviral transduction (vs. pMSCV empty vector, EV). Cells infected with two different titers are shown. **(C)** Visualization of SDE2-APEX2-mediated proximity biotin labeling of endogenous proteins by streptavidin-horseradish peroxidase (HRP) Western blotting. Negative controls in which BP, $H_2O_2$, or SDE2-APEX2 were omitted are shown. Asterisks indicate endogenous biotinylated proteins known to migrate near 130, 75, and 72 kD.

Figure S2

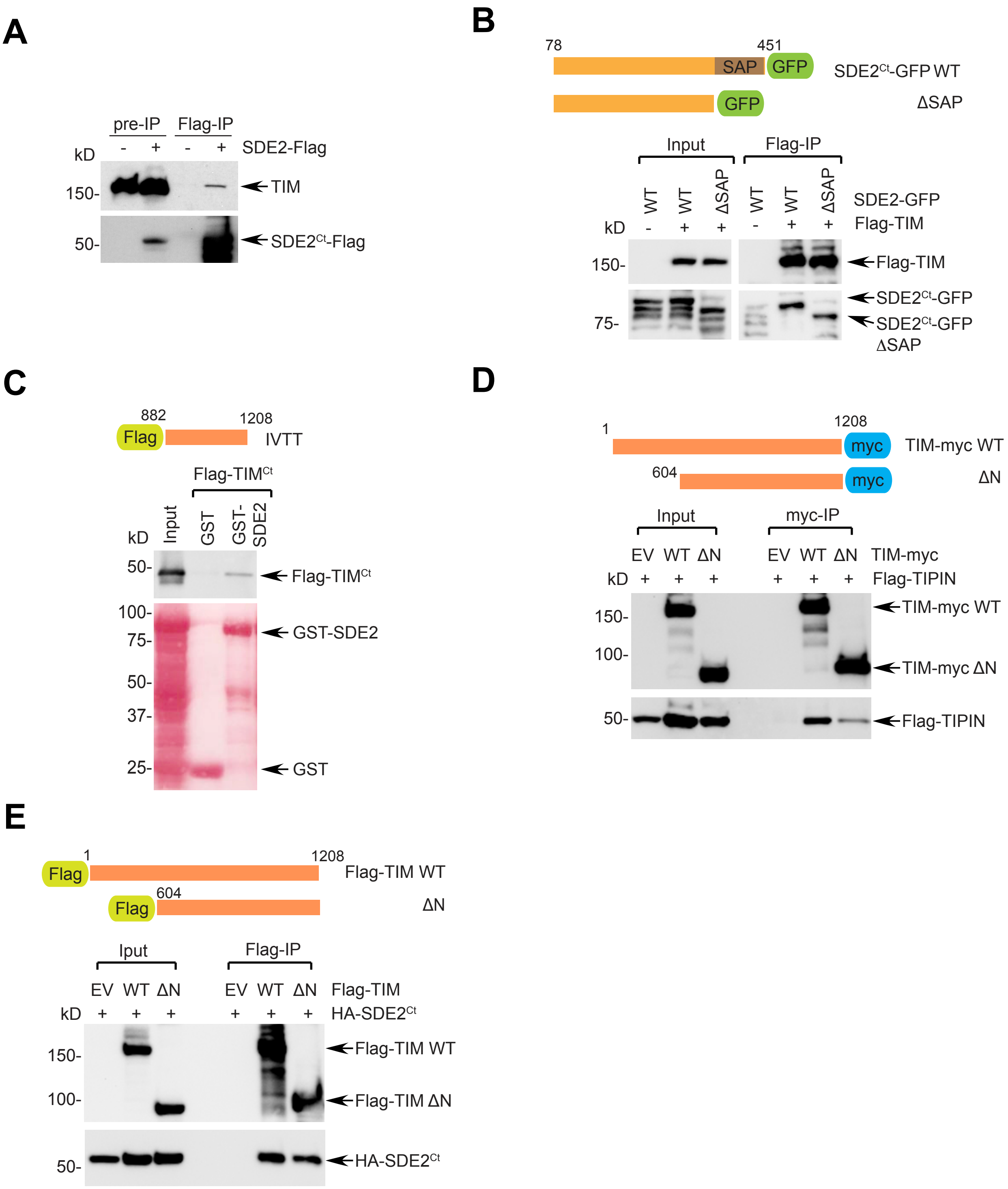
A
pre-IP
Flag-IP
kD
- + - +
SDE2-Flag
150-
TIM
50-
SDE2Ct-Flag
B
78
451
SAP
GFP
SDE2Ct-GFP WT
GFP
ΔSAP
Input
Flag-IP
WT WT ΔSAP WT WT ΔSAP
SDE2-GFP
kD
- + + - + +
Flag-TIM
150-
Flag-TIM
75-
SDE2Ct-GFP
SDE2Ct-GFP ΔSAP
C
882
1208
Flag
IVTT
Flag-TIMCt
kD
Input
GST
GST-SDE2
50-
Flag-TIMCt
100-
75-
GST-SDE2
50-
37-
25-
GST
D
1
1208
myc
TIM-myc WT
604
myc
ΔN
Input
myc-IP
EV WT ΔN EV WT ΔN
TIM-myc
kD
+ + + + + +
Flag-TIPIN
150-
TIM-myc WT
100-
TIM-myc ΔN
50-
Flag-TIPIN
E
1
1208
Flag
Flag-TIM WT
604
Flag
ΔN
Iput
Flag-IP
EV WT ΔN EV WT ΔN
Flag-TIM
kD
+ + + + + +
HA-SDE2Ct
150-
Flag-TIM WT
100-
Flag-TIM ΔN
50-
HA-SDE2Ct

**Supplementary Figure S2 (related to Figure 2).**

**(A)** Anti-Flag co-immunoprecipitation (IP) of SDE2-Flag with endogenous TIM in U2OS cells. (-: empty vector). **(B)** Anti-Flag co-IP of Flag-TIM with SDE2-GFP WT, or ΔSAP (Δ385-451) mutant in 293T cells. Schematic of the SDE2-GFP construct variants is shown above. **(C)** GST pull-down of *in vitro* transcribed and translated (IVTT) $TIM^{Ct}$ (aa882-1208) with purified GST or GST-SDE2. **(D)** Anti-myc co-IP of TIM-myc WT or Δ1-603 (ΔN) mutant with Flag-TIPIN in 293T cells. Schematic of the TIM-myc construct variants is shown above. **(E)** Anti-Flag co-IP of Flag-TIM WT or ΔN mutant with HA-$SDE2^{Ct}$ in 293T cells.

Figure S3

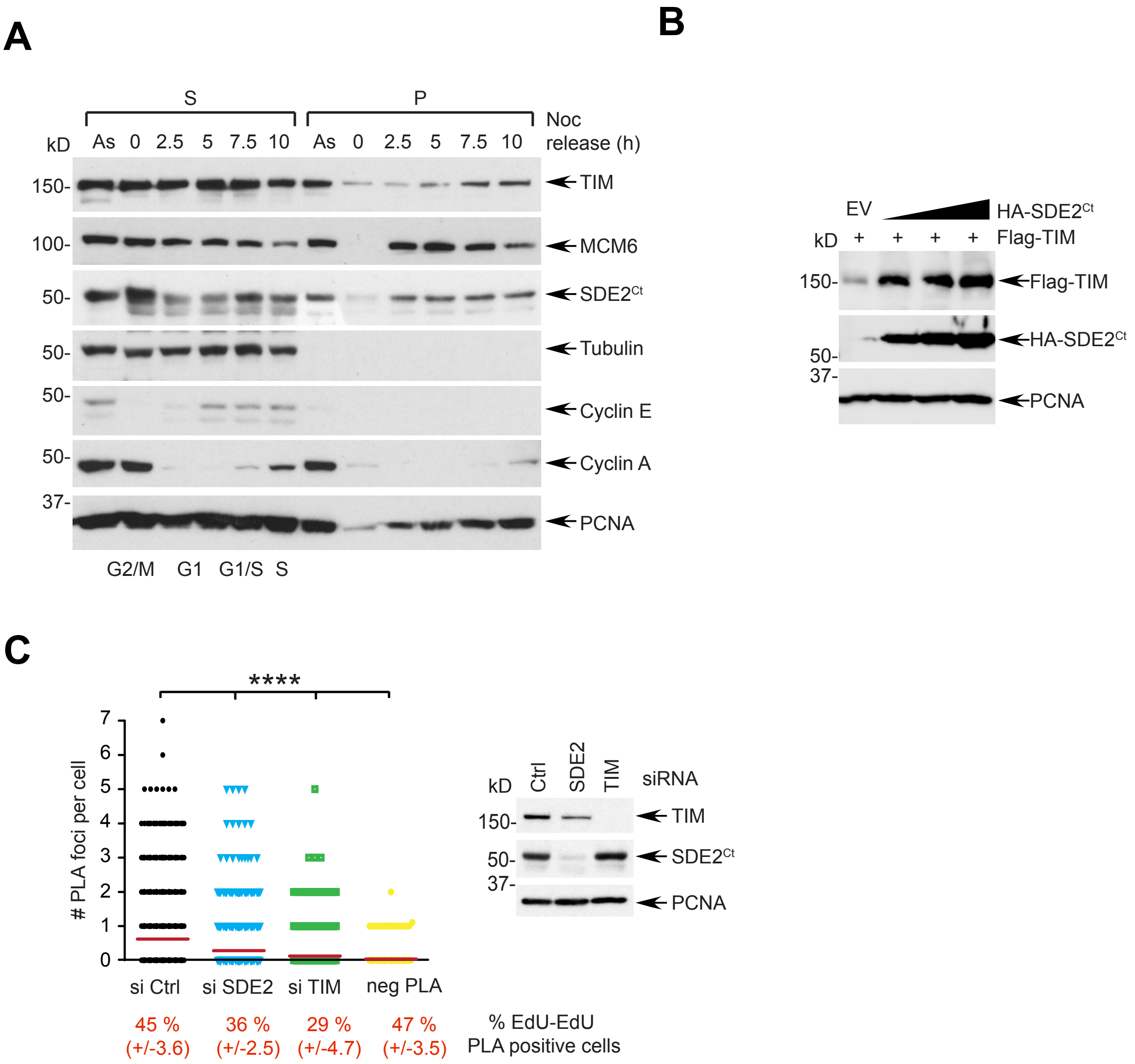
A
S
P
Noc release (h)
kD
As 0 2.5 5 7.5 10 As 0 2.5 5 7.5 10
150-
TIM
100-
MCM6
50-
SDE2Ct
50-
Tubulin
50-
Cyclin E
50-
Cyclin A
37-
PCNA
G2/M G1 G1/S S
B
EV
HA-SDE2Ct
kD
+ + + +
Flag-TIM
150-
Flag-TIM
HA-SDE2Ct
50-
37-
PCNA
C
****
### PLA foci per cell
7
6
5
4
3
2
1
0
si Ctrl
si SDE2
si TIM
neg PLA
45 % (+/-3.6)
36 % (+/-2.5)
29 % (+/-4.7)
47 % (+/-3.5)
% EdU-EdU PLA positive cells
kD
Ctrl
SDE2
TIM
siRNA
150-
TIM
50-
SDE2Ct
37-
PCNA

**Supplementary Figure S3 (related to Figure 3).**

**(A)** Cell cycle-dependent changes of TIM and $SDE2^{Ct}$ levels in both S and P fractions. U2OS cells were synchronized at the G2/M boundary with 100 ng/mL nocodazole and released into fresh medium to let cells traverse from G1 to S phase. Cells were harvested at the indicated times and fractionated into S and P fractions, followed by WB analysis. **(B)** Elevation of cellular Flag-TIM levels in 293T cells in response to increasing amount of HA-$SDE2^{Ct}$. **(C)** Left: quantification of TIM:EdU PLA foci numbers from PLA-positive cells. The percentage of EdU:EdU PLA positive cells is shown. The number of foci per positive cells was pooled from three independent experiments (*****P*<0.0001, Mann-Whitney test). Right: WB analysis to confirm knockdown efficiency.

Figure S4

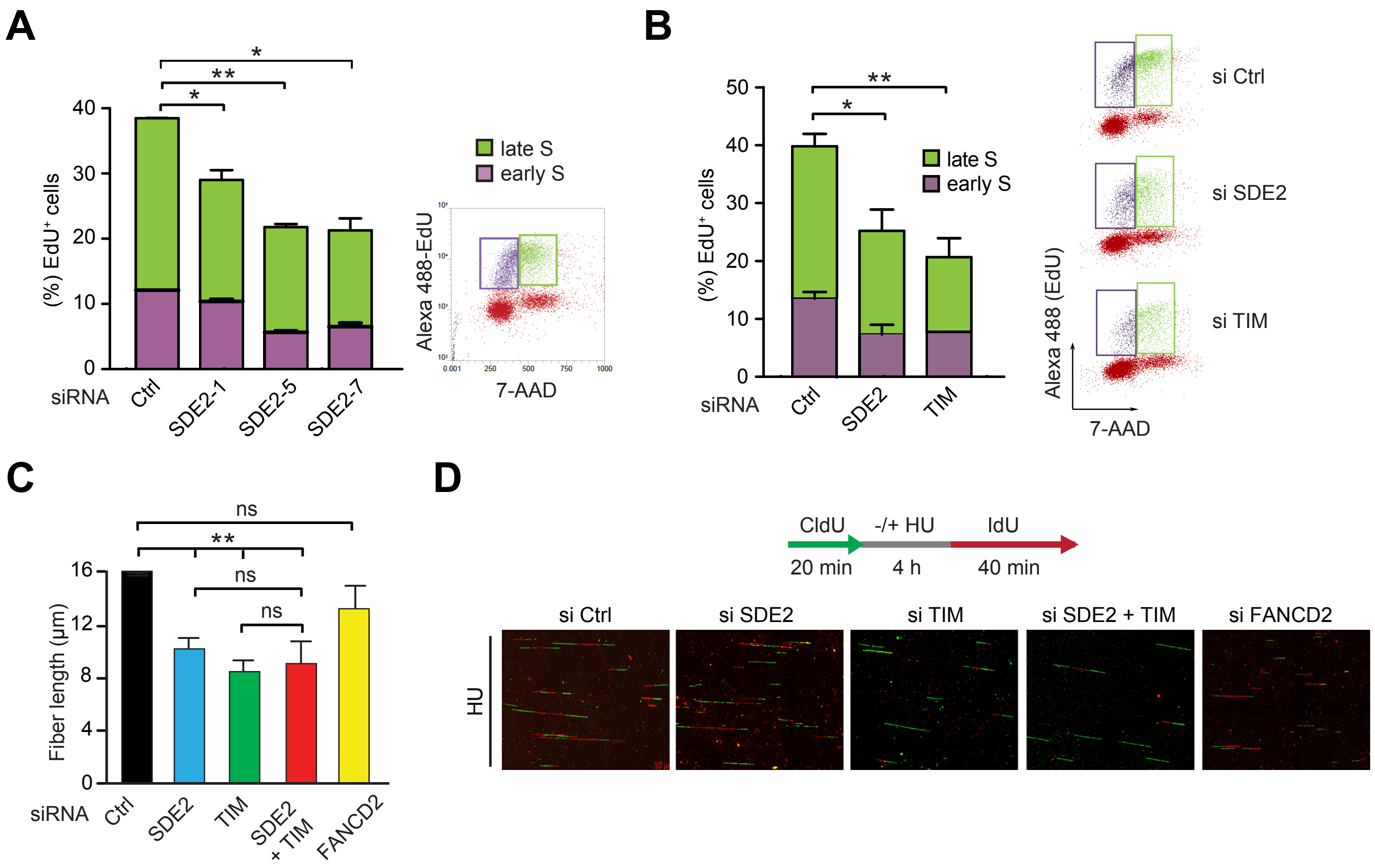
A
late S
early S
(%) EdU⁺ cells
siRNA
Ctrl
SDE2-1
SDE2-5
SDE2-7
Alexa 488-EdU
7-AAD
B
(%) EdU⁺ cells
siRNA
Ctrl
SDE2
TIM
si Ctrl
si SDE2
si TIM
Alexa 488 (EdU)
7-AAD
C
Fiber length (μm)
siRNA
Ctrl
SDE2
TIM
SDE2 + TIM
FANCD2
ns
D
CldU
-/+ HU
IdU
20 min
4 h
40 min
si Ctrl
si SDE2
si TIM
si SDE2 + TIM
si FANCD2
HU

**Supplementary Figure S4 (related to Figure 4).**

**(A)** Left: EdU cell cycle analysis of U2OS cells transfected with independent SDE2 siRNA oligos. (n=2, mean ± SD, ***P*<0.01, **P*<0.05, Student's t-test, for the comparison of late S phase cells). Right: a representative FACS plot defining early and late S phases from EdU-positive cells **(B)** EdU cell cycle analysis of U2OS cells transfected with the indicated siRNAs (n=2, mean ± SD, ***P*<0.01, **P*<0.05, Student's t-test, for the comparison of late S phase cells). Representative FACS plots showing the distribution and percentage of EdU-positive cells in early and late S phases are shown. **(C)** Mean IdU DNA fiber track lengths from U2OS cells transfected with indicated siRNAs (n=3, mean ± SD, ***P*<0.01, Student's t-test, ns, not significant). **(D)** Representative images of DNA fiber tracks after HU-mediated fork stalling in U2OS cells transfected with the indicated siRNAs. Increase of green track-only DNA fibers represents increased fork stalling.

### Figure S5

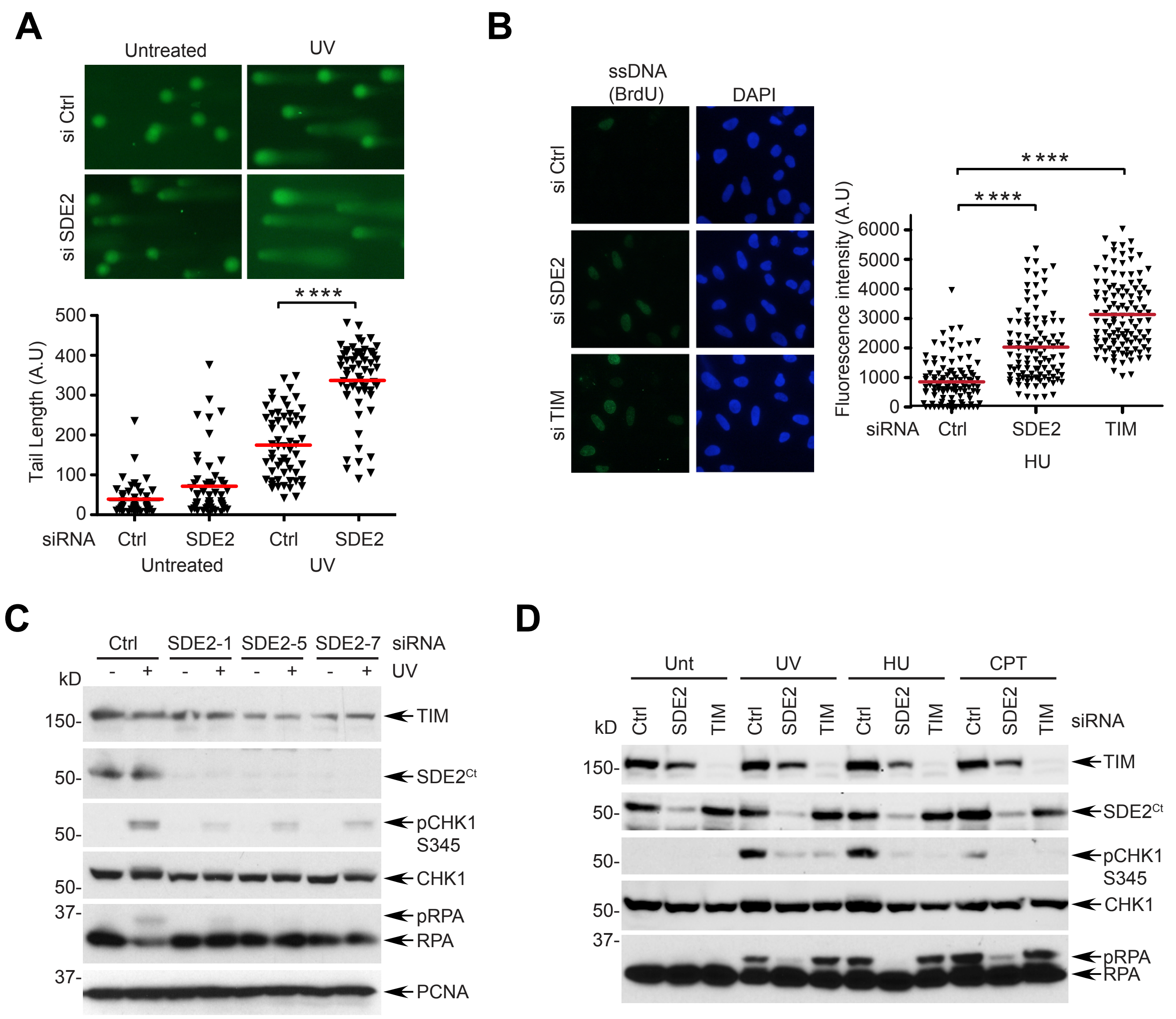
A
Untreated
UV
si Ctrl
si SDE2
Tail Length (A.U)
500
400
300
200
100
0
****
siRNA Ctrl SDE2 Ctrl SDE2
Untreated UV
B
ssDNA (BrdU)
DAPI
si Ctrl
si SDE2
si TIM
Fluorescence intensity (A.U)
6000
5000
4000
3000
2000
1000
0
****
****
siRNA Ctrl SDE2 TIM
HU
C
Ctrl SDE2-1 SDE2-5 SDE2-7 siRNA
- + - + - + - + UV
kD
150-
50-
50-
50-
37-
37-
TIM
SDE2Ct
pCHK1 S345
CHK1
pRPA
RPA
PCNA
D
Unt UV HU CPT
Ctrl SDE2 TIM Ctrl SDE2 TIM Ctrl SDE2 TIM Ctrl SDE2 TIM siRNA
kD
150-
50-
50-
50-
37-
TIM
SDE2Ct
pCHK1 S345
CHK1
pRPA
RPA

**Supplementary Figure S5 (related to Figure 5).**

**(A)** Top: representative images of DNA comets from siRNA-transfected U2OS cells 4 h after irradiation with 40 J/m$^2$ UVC. Bottom: quantification of DNA comet tail lengths from two independent experiments. Red bars represent median (>50 nuclei per condition, ****$P$<0.0001, Mann-Whitney test). **(B)** Left: representative images of native BrdU staining after treatment with 2 mM HU for 4 h in U2OS cells transfected with the indicated siRNAs. Cells were pre-treated with 10 μM BrdU for 48 h before irradiation to mark single-stranded DNA. Right: quantification of BrdU staining. Red bars represent median (>100 nuclei per condition, n=2, ****$P$<0.0001, Mann-Whitney test). **(C)** CHK1 phosphorylation at S345 4 h after irradiation with 40 J/m$^2$ UVC in U2OS cells transfected with independent SDE2 siRNA oligos. **(D)** CHK1 phosphorylation after the indicated types of DNA damages (2 mM HU for 4 h, 4 h after 40 J/m$^2$ UVC irradiation, or 100 nM camptothecin (CPT) for 4 h) in U2OS cells transfected with the indicated siRNAs.

### Figure S6

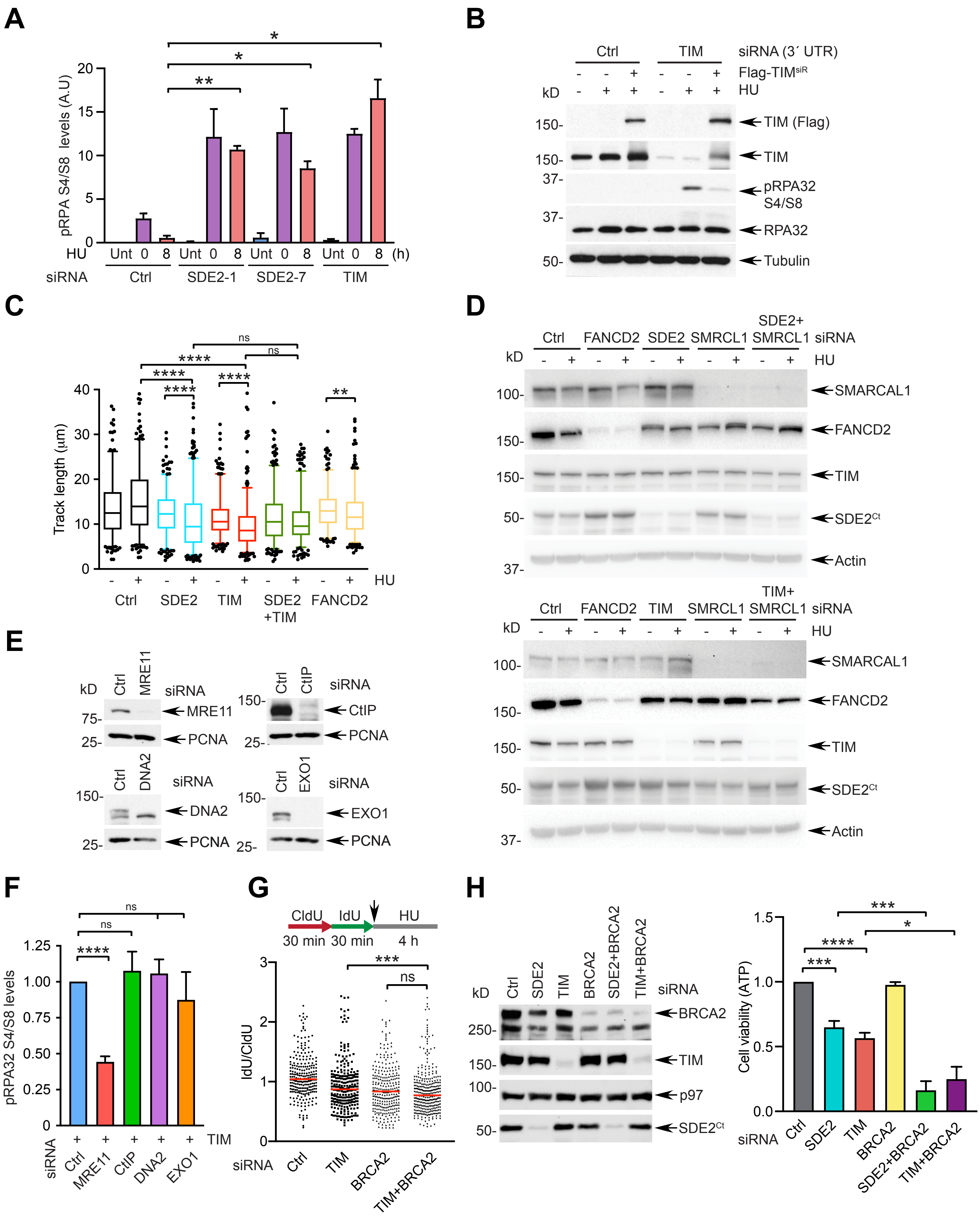

A
pRPA S4/S8 levels (A.U)
HU Unt 0 8 Unt 0 8 Unt 0 8 Unt 0 8 (h)
siRNA Ctrl SDE2-1 SDE2-7 TIM
B
siRNA (3´ UTR) Ctrl TIM
Flag-TIMsiR
HU
TIM (Flag)
TIM
pRPA32 S4/S8
RPA32
Tubulin
C
Track length (µm)
HU
Ctrl SDE2 TIM SDE2 +TIM FANCD2
D
Ctrl FANCD2 SDE2 SMRCL1 SDE2+ SMRCL1 siRNA
HU
SMARCAL1
FANCD2
TIM
SDE2Ct
Actin
Ctrl FANCD2 TIM SMRCL1 TIM+ SMRCL1 siRNA
E
MRE11
CtIP
DNA2
EXO1
PCNA
F
pRPA32 S4/S8 levels
TIM
siRNA Ctrl MRE11 CtIP DNA2 EXO1
G
CldU 30 min IdU 30 min HU 4 h
IdU/CldU
siRNA Ctrl TIM BRCA2 TIM+BRCA2
H
siRNA Ctrl SDE2 TIM BRCA2 SDE2+BRCA2 TIM+BRCA2
BRCA2
TIM
p97
SDE2Ct
Cell viability (ATP)

**Supplementary Figure S6 (related to Figure 6).**

**(A)** Quantification of pRPA32 S4/S8 levels in siRNA-transfected U2OS cells recovered from the overnight treatment of 250 μM HU (n=2, mean ± SEM, **P*<0.05, ***P*<0.01, Student's t-test). **(B)** Rescue of the elevated pRPA32 S4/S8 phenotype. U2OS cells were serially transfected with siRNA TIM that targets the 3´ UTR of *TIM* mRNA and Flag-TIM encoding plasmid. Cells were treated with 250 μM HU for 20 h and recovered for 4 h in fresh medium to assess pRPA32 S4/S8 levels. **(C)** Distributions of the CldU track length from siRNA-transfected U2OS cells untreated (unt) or treated with 2 mM HU for 4 h. FANCD2 knockdown serves as a positive control showing impaired fork protection. The median value of at least 300 tracks per experimental condition is indicated (n=3, *****P*<0.0001, ***P*<0.01, ns, not significant, Mann-Whitney test). **(D)** WB analysis to confirm knockdown of SDE2 and TIM individually or together with SMARCAL1 in U2OS cells, used for DNA combing analysis. **(E)** WB analysis to confirm siRNA knockdown of individual nucleases. **(F)** Quantification of pRPA32 S4/S8 levels in U2OS cells co-depleted of TIM and individual nucleases upon recovery from overnight treatment of 250 μM HU (n=3, mean ± SD, *****P*<0.0001, Student's t-test). **(G)** Dot plot of DNA fiber IdU/CldU track length ratios from U2OS cells either knocked-down with TIM only or TIM and BRCA2 together. A representative plot is shown from three independent experiments (n=3, ****P*<0.001, Mann-Whitney). **(H)** Left: WB analysis to confirm the knockdown of SDE2 and TIM individually or in combination with BRCA2. Right: luminescence-based ATP viability assay of U2OS cells transfected with the indicated siRNAs. Measurement was performed 7 days after transfection (n=3, mean ± SD, *****P*<0.0001, ****P*<0.001, **P*<0.05, Student's t-test).

### Figure S7

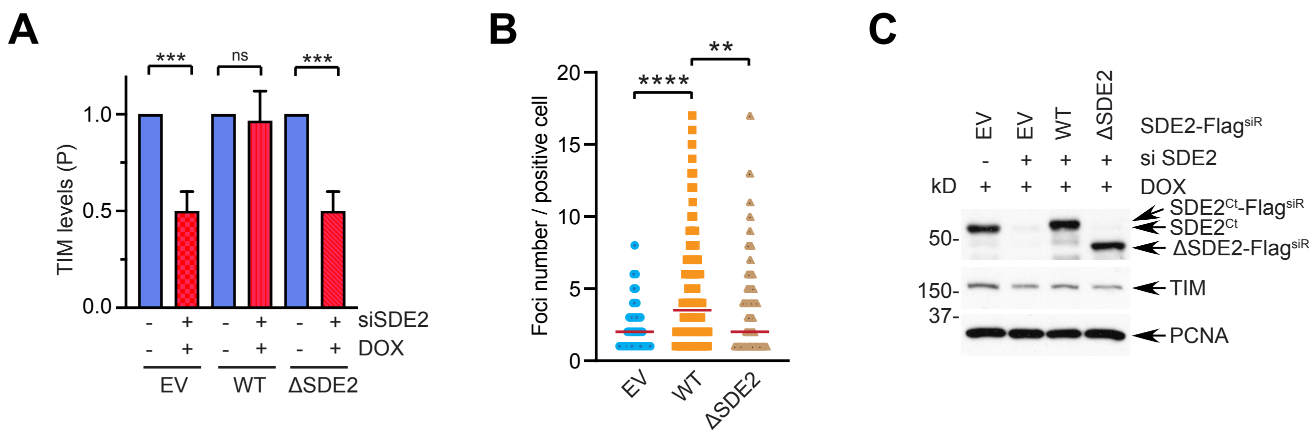

A
***
ns
***
TIM levels (P)
1.0
0.5
0.0
siSDE2
DOX
EV
WT
ΔSDE2
B
Foci number / positive cell
20
15
10
5
0
****
**
EV
WT
ΔSDE2
C
SDE2-FlagsiR
si SDE2
DOX
kD
50-
150-
37-
SDE2Ct-FlagsiR
SDE2Ct
ΔSDE2-FlagsiR
TIM
PCNA

**Supplementary Figure S7 (related to Figure 7).**

**(A)** Quantification of TIM levels in the P fraction from subcellular-fractionated Retro-X SDE2 WT or ΔSDE2 cells following SDE2 siRNA transfection and doxycycline (dox) induction (n=3, mean ± SD, ***$P$<0.001, Student's t-test). **(B)** Quantification of TIM PLA:EdU foci numbers from the PLA-positive cells. Red bars represent median (n=3 pooled from three independent experiments, ****$P$<0.0001, **$P$<0.01, Mann-Whitney test). **(C)** WB analysis to confirm the reconstitution of Flag-tagged SDE2 WT or ΔSDE2 mutant by dox induction following siRNA transfection in Retro-X U2OS cells used for the DNA combing analyses in Figures 7D and 7E.
